# Functional Divergence of GIGANTEA paralogs in *Petunia x hybrida* Reveals Uncoupling of Vegetative Growth and Flowering Time

**DOI:** 10.64898/2026.01.20.700548

**Authors:** Claudio Brandoli, Marcos Egea-Cortines, Julia Weiss

**Author notes:** Corresponding author: Julia Weiss, Claudio Brandoli, Marcos Egea-Cortines.

## Abstract

The circadian clock gene *GIGANTEA* (GI) coordinates vegetative growth and floral transition in plants, yet the roles of its paralogs in Solanaceae remain unclear. In *Petunia × hybrida*, the GI locus is duplicated as *PhGI1* and *PhGI2*. We investigated whether this duplication uncouples vegetative growth from flowering time. Using RNA interference and CRISPR/Cas9 mutagenesis, we dissected the function of *PhGI2*. RNAi silencing of *PhGI2* produced complex phenotypes, including enhanced vegetative growth, early floral abortion, and delayed flowering, raising questions about cosuppression effects. To resolve this, we generated three independent CRISPR alleles of *PhGI2*. These mutants exhibited extreme late flowering without changes in vegetative architecture or scent emission, demonstrating that *PhGI2* specifically controls floral transition. Expression analyses revealed that *PhGI2* is required to maintain rhythmicity and amplitude of *PhGI1* and core clock genes, including *PhTOC1* and *PhLHY*. Our findings uncover subfunctionalization of *GI* paralogs. Whilst *PhGI1* represses vegetative growth and influences floral development, *PhGI2* promotes flowering and coordinates circadian transcription. This genetic separation of vegetative growth and flowering time provides a framework for manipulating crop architecture and phenology.

**Highlight:** Duplication of GIGANTEA in petunia uncouples vegetative growth from flowering time through paralog subfunctionalization

## Introduction

The coordination of plant development with the environment is partly orchestrated by the plant circadian clock, which regulates processes such as flowering time, stomatal opening, cell proliferation and expansion, and floral scent emission (Liu *et al*., 2001; Nusinow *et al*., 2011; Fenske *et al*., 2015; Fung-Uceda *et al*., 2018). A common set of clock-associated genes, found in the picoeukaryote *Ostreococcus* and conserved in plants, include a MYB transcription factor *LATE ELONGATED HYPOCOTYL (LH*Y), a *PSEUDO RESPOSE REGULATOR (TOC1)*, and a blue light receptor similar to ZEITLUPE (ZTL) (Corellou *et al*., 2009; Bouget *et al*., 2011). Early in the evolution of land plants, the genetic architecture of the plant circadian clock became more complex, incorporating additional clock components and interlocking feedback loops (McClung, 2006; Staiger *et al*., 2013). One such gene is *GIGANTEA* (*GI*) which is found in *Marchantia polymorpha* and some charophytes but is absent in *Physcomitrium patens* or *Selaginella moellendorffii* (Linde *et al*., 2017)*. AtGI* was originally identified in Arabidopsis as a mutant with delayed flowering and increased vegetative size (Rédei, 1962; Fowler *et al*., 1999).

The biological functions of *AtGI* in flowering time occur partly via its role in the circadian clock. Loss of function of *AtGI* causes a shorter circadian rhythm with dampened rhythmic expression of clock genes such as *AtLHY* under normal and free running conditions (Mizoguchi *et al*., 2005; Sawa *et al*., 2007). It is also a positive regulator of the flowering time gene *AtFT* in Arabidopsis (Sawa and Kay, 2011). This function is conserved in *Late Bloomer1*, the pea ortholog of *GI* (Hecht *et al*., 2011) *Marchantia polymorpha* (Kubota *et al*., 2014), soybean, or poplar (Watanabe *et al*., 2011; Dong *et al*., 2022; Wang *et al*., 2023; Alique *et al*., 2024). It is also responsible for seasonal adaptation in barley (Zakhrabekova *et al*., 2012). Notably, the two soybean *E2* maturity genes are *GI* orthologs that redundantly regulate photoperiodic flowering and yield (Hayama *et al*., 2003; Wang *et al*., 2023). Thus, *GI* has a dual biological function in controlling both vegetative growth and flowering time.

The *GIGANTEA* (GI) locus is duplicated in several plant lineages, including members of the Solanaceae, with reported copy numbers ranging from two in *Petunia × hybrida* to three in *P. integrifolia* and *Solanum lycopersicum* (Bombarely *et al*., 2016). However, the specific roles of individual *GI* paralogs remain largely unexplored. Functional analysis of *PhGI1* in *P. hybrida* through gene silencing revealed a complex phenotype, including enhanced vegetative growth, ectopic floral development, early floral senescence, and a slight reduction in flower size. Floral scent emission was only mildly affected (Brandoli *et al*., 2020). Interestingly, silencing *PhGI1* also led to a significant decrease in the expression of *PhGI2*, raising the question of whether the observed phenotypic changes are due solely to the loss of *PhGI1*, or whether downregulation of *PhGI2* also contributes. Moreover, it remains unclear whether the effect on *PhGI2* expression is an off-target consequence of the silencing construct, or whether *PhGI1* and *PhGI2* are involved in a regulatory feedback mechanism governing each other’s expression. However, silencing of *PhGI1* does not cause a delayed flowering phenotype suggesting a possible functional divergence of *GI* in Petunia.

Here, we present a comprehensive functional analysis of the *Petunia x hybrida PhGI2* paralog, using RNAi silencing and CRISPR/Cas9-mediated targeted mutagenesis. Our results show that silencing of *PhGI2* caused a series of phenotypes previously found in *RNAi::PhGI1*. However, *RNAi::PhGI2* also showed delayed flowering, a phenotype not found in *RNAi::PhGI1*. We hypothesized that by using CRISPR alleles of *PhGI2* we would identify the specific functions of *PhGI2*. Our results show that *PhGI1* and *PhGI2* have undergone subfunctionalization and neofunctionalization. *PhGI2* retained the ancestral function of promoting floral transition and is required for canonic circadian clock gene expression, while *PhGI1* retained the ancestral function of repressing vegetative growth and acquired new roles in flower development.

## Material and Methods

### Design of the RNAi and Crispr/Cas9 constructs

We obtained the *PhGI2* coding region from the genome sequence of *Petunia x hybrida* W115. The coding gene model corresponds to Peaxi162Scf00160g01744.1. We developed a hairpin construct that would discriminate *PhGI2* from *PhGI1* for the vector construction targeting the 3’UTR of *PhGI2.* Site-specific primers (Supplementary Table S1) with the attB1 and attB2 sites for Gateway® recombination, were used to PCR-amplify a DNA fragment of 208 bp. To obtain a hairpin-like structure, the *PhGI2* fragment was first recombined into the entry vector pDONR201 (Invitrogen) and then into the destination vector pHELLSGATE 12 (Helliwell and Waterhouse, 2003).

The Crispr/Cas9 guide targeting *PhGI2* was selected using the web tool CHOPCHOP (Labun *et al*., 2019) which offers analysis of the target genome of *Petunia x hybrida*. A vector VB211218-1015kpn pPBV[CRISPR]-Neo/Kana-zCas9-AtU6-26>{PhGI2_exon4} of 15224 bp was build and purchased, with a guide RNA targeting exon 4 of the gene *PhGi2* (ACTGCCTTCAACTCCTAGGT). Integrity of all constructs were confirmed through PCR amplification and visualization on 1% agarose gel.

### Plant material, transformation and sampling

Seeds of *Petunia x hybrida* of the double haploid variety ’Mitchell W115’ were collected. *In vitro* germinated plants were used as the wildtype controls and as the source of explants for plant transformation as described (Manchado-Rojo *et al*., 2014). The disarmed *Agrobacterium tumefaciens* strain EHA105 was used for transformation as described previously. Lines of T0 and T1 generation transformed with Crispr/Cas9 constructs, were confirmed through PCR detection of the *ZmCAS9*gene.

Four independent T0 lines *RNAi*::*PhGI2* were selected for further studies. The T1 generation of wildtype plants as well as silenced lines of *PhGI2*, were grown in a growth chamber under controlled conditions of 16 hours of light/8 hours of darkness (16:8 LD), luminous intensity of 250 μE m ^-2^ s ^-1^ and a constant temperature of 26 ± 1° C. The T2 generations were cultured in a greenhouse under natural long-day conditions.

The T1 generation of seven independent T0 Crispr/Cas9-*PhGI2* lines were further phenotyped. Plants were grown in a greenhouse a photoperiod ranging from 9.5 – 12.5 h of day light. We obtained three independent alleles that were used further.

### Circadian sampling

Samples were taken every three (*RNAi*::*PhGI2)* to four (Crispr/Cas9-*PhGi2)* hours from wildtype plants during 24 hours and one or two plants from independent silenced lines as well as two early flowering control plants and two late flowering mutant lines homozygous for the Crispr/Cas9-*PhGI2* alleles. The collected tissues were immediately frozen in liquid nitrogen and stored at –80°C until further analysis.

Under growth chamber conditions, ZEITGEBER Time 0 (ZT0) was considered as the time when the light was turned on. Under natural greenhouse conditions, sampling for expression analysis was conducted at sunrise under 12 hours day light condition with ZEITGEBER Time 0 (ZT0) coinciding with sunrise. We performed experiments under free running conditions using WT plants that had grown for several weeks under long day conditions and were transferred to continuous dark.

### Phenotypic analysis

For each selected RNAi::*PhGI2* T1 line, T2 plants were propagated after self-pollination. At least three T2 plants were characterized for each line to analyse the phenotypes associated with *GI2* RNA interference. We analysed vegetative growth including plant height, internode length, number of leaves to first flower, number of axillary stems, leaf length and width. We quantified number of flower buds and fully developed flowers, corolla diameter, tube and petiole length.

T1 populations of 30-40 individuals from Crispr/Cas9-*PhGI2* lines were grown. Three lines (4,12 and 19) showed a clear segregation into early flowering, mid flowering and late flowering phenotypes and were further phenotyped in detail. The *GI2* region targeted by the Crispr/Cas9 construct was amplified and sequenced for three late flowering plants of each line. We analysed vegetative growth including stem length, branch number, internode length, leaf length and width. We measured flower number, days to first flower, corolla width and tube length.

### Volatile organic compound analysis

The analysis of volatile organic emission compound (VOC) was performed sampling three flowers from wildtype plants and two plants of two silenced lines after 2-3 days after anthesis as described in (Manchado-Rojo *et al*., 2012). For each of the three Crispr/CAS9-*PhGI2* lines, we analysed VOC emission of four flowers from early flowering control plants and late flowering mutant plants during a period of 4 hours, starting at sunrise.

### Analysis of circadian gene expression

We used phenol:chloroform to isolate total RNA from leaves (Box *et al*., 2011) followed by spectrophotometric quantification (NanoDrop2000). Equal amounts of RNA were used to synthesize cDNA according to the manufacturer’s instructions (Maxima First Strand cDNA Synthesis Kit for RT-qPCR, with dsDNase, https://www.thermofischer.com/, catalog number: K1641). Three biological samples in form of young leaves and two technical replicas were analysed for each sample in quantitative PCR analysis. The gene *ACTIN 11* (*ACT*), was used as reference gene for relative gene expression quantification, after selection as valuable housekeeping gene for Petunia leaves and petals under circadian conditions (Terry *et al*., 2019a). All the primers used for *PhGI2, PhGI1* and other clock genes were designed using pcrEfficiency software (Mallona *et al*., 2011) (Supplementary Table S1).

### Data analysis

The relative gene expression of the circadian genes, relative to the reference gene ACT, was calculated applying the comparative CT method (Schmittgen and Livak, 2008) as well as using group-wise comparison with the REST Program (Pfaffl *et al*., 2002). Periodicity and significance were evaluated using JTK_CYCLE and Lomb–Scargle (LS) as implemented in MetaCycle (R version 4.3.2) (Glynn *et al*., 2006; Hughes *et al*., 2010; Wu *et al*., 2016). P-values were adjusted with the Benjamini-Hochberg FDR (Benjamini and Hochberg, 1995).

Significance differences among data were determined based on Fisheŕs F-test and Student’s T-Test after data transformation to fit to a normal distribution. Volatile organic compound profiles were analyzed using the R-package GCprofileMaker (Perez-Sanz *et al*., 2021).

## 3. Results

### Generation of RNAi and CRISPR/Cas9 *PhGI2* Lines

To investigate the biological role of *PhGI2*, we generated and analysed two types of loss-of-function lines: four independent RNA interference (RNAi) lines (RNAi::*PhGI2.4.2*, *4.4, 6.1*, and *6.2*) and three CRISPR/Cas9-induced mutant alleles (*PhGI2.12*, *PhGI2.19*, and *PhGI2.4*). Initial RNAi constructs targeting *PhGI1 3’-UTR* resulted in ∼50% suppression of *PhGI2* (Brandoli *et al*., 2020), suggesting either cosuppression effects (Angenent *et al*., 1994) or potential transcriptional regulation of *PhGI2* by *PhGI1*. To resolve this, we designed RNAi fragments that specifically targeted *PhGI2 3’-UTR*, avoiding sequence overlap with *PhGI1*.

To further isolate *PhGI2*-specific functions, we developed CRISPR/Cas9 alleles using guide RNAs targeting unique sequences in exon 4 of *PhGI2*. We characterized three alleles. *PhGI2.19* had a three-base pair in-frame deletion removing the highly conserved Proline 215 (P215) (Supplementary Fig. S1). *PhGI2.4* had a single base insertion causing a frameshift at position 216, generating five altered amino acids followed by a premature stop codon. Finally, *PhGI2.12* had a four-base deletion leading to an 11-amino acid frameshift starting at position 215, also followed by a stop codon (Fig. 1A-C).

**Fig. 1.**
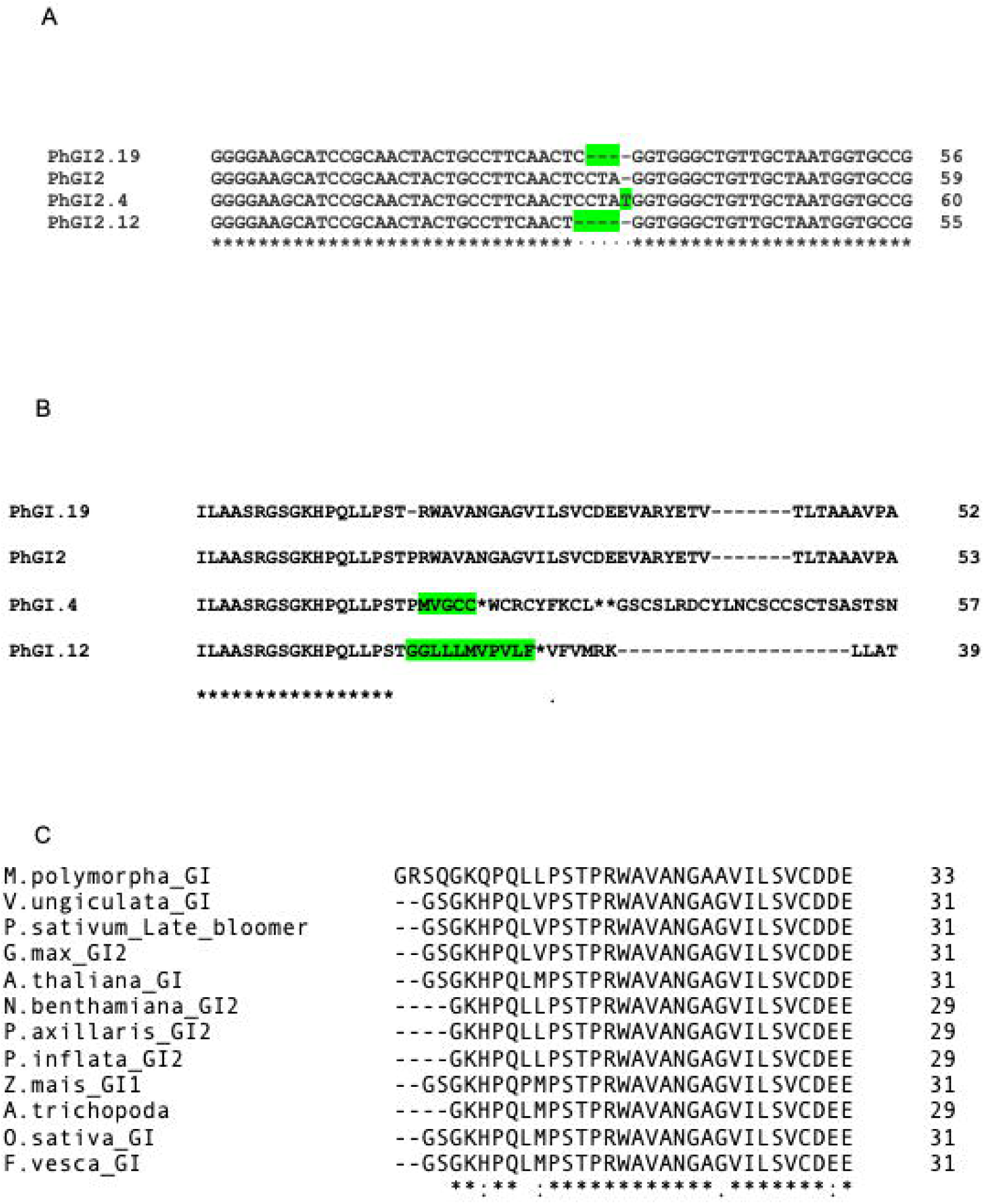
Mutations in *PhGI2* obtained by CRISPR/Cas9. (A) Sequence alignment of the *GI2* exon4 region. Deletions and insertions are marked in green. (B) Positions in the protein mutated in *PhGI2.4*, *PhGI2.12* and *PhGI.19.* Amino acids marked in green correspond to the new sequence resulting from the frame shift. (C) Conserved region in GI proteins across the plant kingdom.

The predicted wild-type PhGI2 protein is 1162 amino acids, consistent in length with GI homologs in *Oryza sativa* (1160), *Arabidopsis thaliana* (1173), and *Marchantia polymorpha* (1187). In contrast, *PhGI2.12* and *PhGI2.4* encode truncated proteins of only 215–216 amino acids. These premature truncations likely disrupt protein function.

Similar truncations in *Arabidopsis GI* alleles, such as *gi-2* (aa144), *gi-6* (aa492), and *gi-3* (aa963) result in late flowering (Araki and Komeda, 1993; Fowler *et al*., 1999). Notably, the deleted P215 in *PhGI2.19* is evolutionarily conserved among GI proteins across plant species (Fig. 1C), supporting the hypothesis that these alleles represent loss-of-function mutations.

Collectively, these silenced and mutant lines enabled us to investigate the role of *PhGI2* within the context of Petunia circadian regulation.

### *PhGI1* and *PhGI2* lose rhythmicity under free-running conditions

We had previously shown that both *PhGI1* and *PhGI2* appear to lose expression under continuous dark (DD) free running conditions (Brandoli *et al*., 2020). These results suggest that *PhGI1* and *PhGI2* do not function as self-sustained oscillators. Nevertheless, we analysed their expression profiles under continuous dark (DD) free-running conditions. JTK_CYCLE and Lomb–Scargle analyses revealed robust rhythmic expression of *PhGI1* and *PhGI2* under LD conditions, whereas no significant rhythmicity was detected under free-running conditions (Supplementary Table S2). This contrasts with *Arabidopsis thaliana*, where *AtGI* maintains rhythmic expression under constant light or darkness (Fowler *et al*., 1999). These results indicate that Petunia *GI* genes require environmental cues to sustain rhythmic expression, revealing a species-specific difference in circadian clock architecture.

### PhGI2 affects the peak expression and rhythmicity of clock genes

We performed a time-series gene expression relative quantification at 3–4 h intervals. Expression patterns of *PhGI2* and *PhGI1* were analysed in the CRISPR line *PhGI2.12* and in RNAi-silenced lines 4.2 (Fig. 2A–D), 4.4, 6.1, and 6.3. In *PhGI2.12*, *PhGI2* peaked at ZT8, coinciding with WT, and its expression level was unchanged. In contrast, *PhGI1* peaked earlier (ZT8) than in WT (ZT12), although overall expression levels were maintained. In *RNAi::PhGI2* lines, the temporal expression pattern was preserved, but transcript levels of both *PhGI1* and *PhGI2* were significantly reduced, with up to an eightfold decrease for *PhGI2* and an approximately 50% reduction for *PhGI1*. This downregulation of *PhGI1* may result from cross-silencing by the RNAi construct or reflect a regulatory role of PhGI2 in *PhGI1* transcription.

**Fig. 2.**
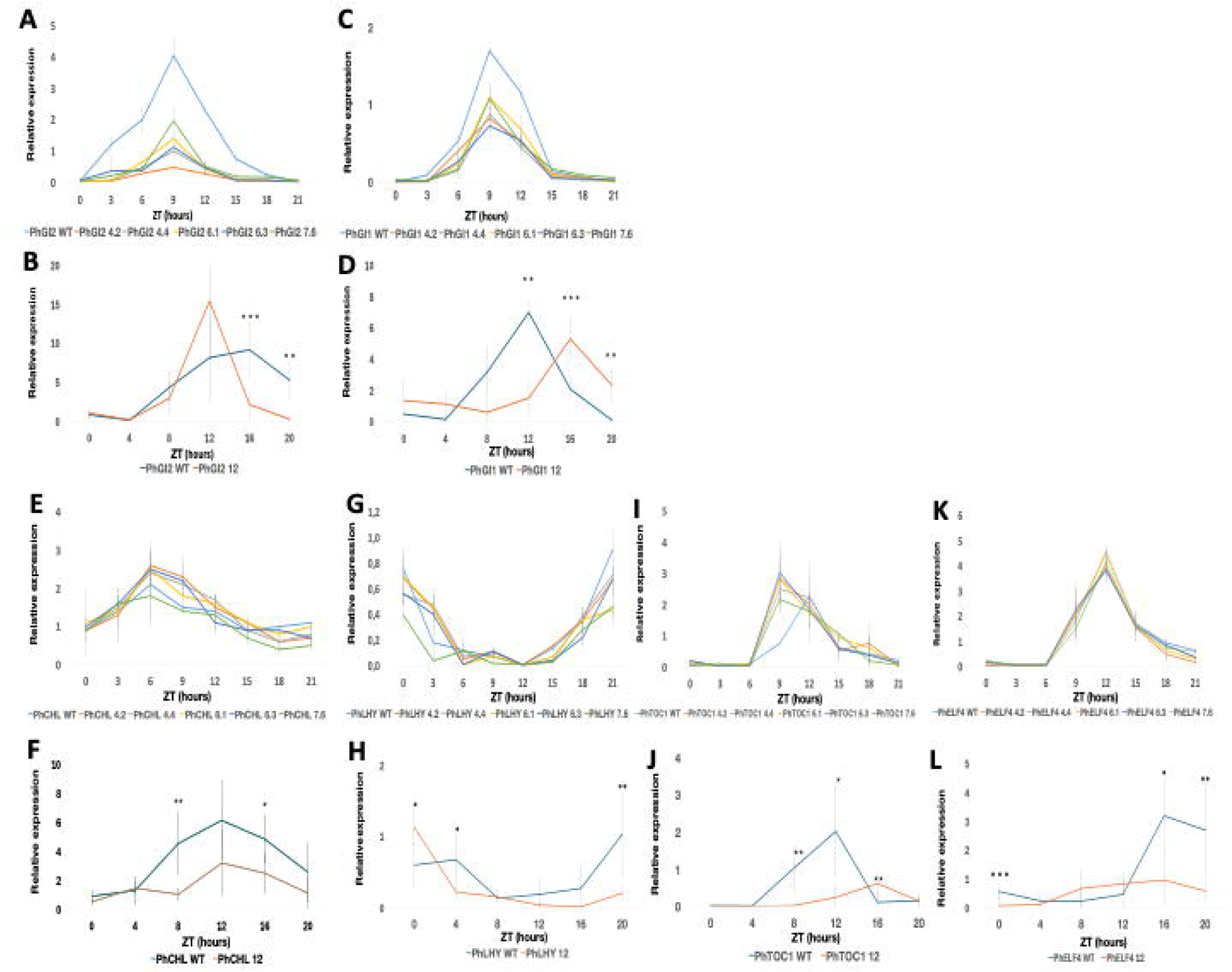
Effect of knocking down and knocking out *PhGI2* on the expression of (A,B) *PhGI2,* (C,D) *PhGI1,* (E,F) *PhCHL*, (G,H) *PhLHY*, (I,J) *PhTOC* and (K,L) *PhELF4* during a 24 hour period in *CRISPR/CAs9 PhGI2* line 12 and in *iRNA::PhGI2* line 4.2, compared to expression in the wild-type. Expression represents the normalized expression NE according to the formula (NE) = 2^-(Ct experimental – Ctn). Three samples were analyzed for each time point and error bars indicate the standard deviation. Asterisks indicate statistical significance between wildtype and iRNA lines with *P < 0.05; **P < 0.01; ***P < 0.001 according to group-wise comparison with to Students T-test.

Silencing of *PhGI2* did not lead to major alterations in the expression profiles of the morning-phased genes *PhLHY*, nor of the evening phased genes *PhCHL*, *PhTOC1* and *PhELF4*, whose temporal patterns remained largely comparable to wild type across the diel cycle (Fig 2. E,G,I,K).

We next investigated the role of *PhGI2.12* in regulating the rhythmic expression of the previously mentioned genes *PhCHL*, *PhLHY*, *PhTOC1*, and *PhELF4* (Fig. 2 F,H,J,L; Supplementary Table S4). All genes displayed robust rhythmic expression in WT plants. Homozygous *PhGI2.12* mutants exhibited advanced expression phases of *PhGI1* and *PhLHY* (4 h) and *PhTOC1* (6 h), accompanied by a loss of rhythmicity in *PhGI1*, *PhCHL*, and *PhELF4*. In addition, expression amplitude was reduced across all analysed genes, with the strongest relative reductions observed for *PhLHY* (66%) and *PhTOC1* (79%). These results demonstrate a strong effect of *PhGI2* on overall circadian gene expression.

### *PhGI2* promotes flowering time but does not affect floral development

We found that *RNAi::PhGI2* lines exhibited significant alterations in flowering time. In the T1 generation, flowering was delayed by approximately 2–3 weeks, while in T2 plants the delay ranged from 1 to 4 weeks (Fig. 3A). Similarly, CRISPR/Cas9 *PhGI2* lines showed a notable delay in flowering time in the T1 generation, with a range of 5–10 weeks between early- and late-flowering individuals, displaying a Mendelian 1:2:1 segregation pattern (Fig. 3B; Supplementary Table S3).

**Figure 3.**
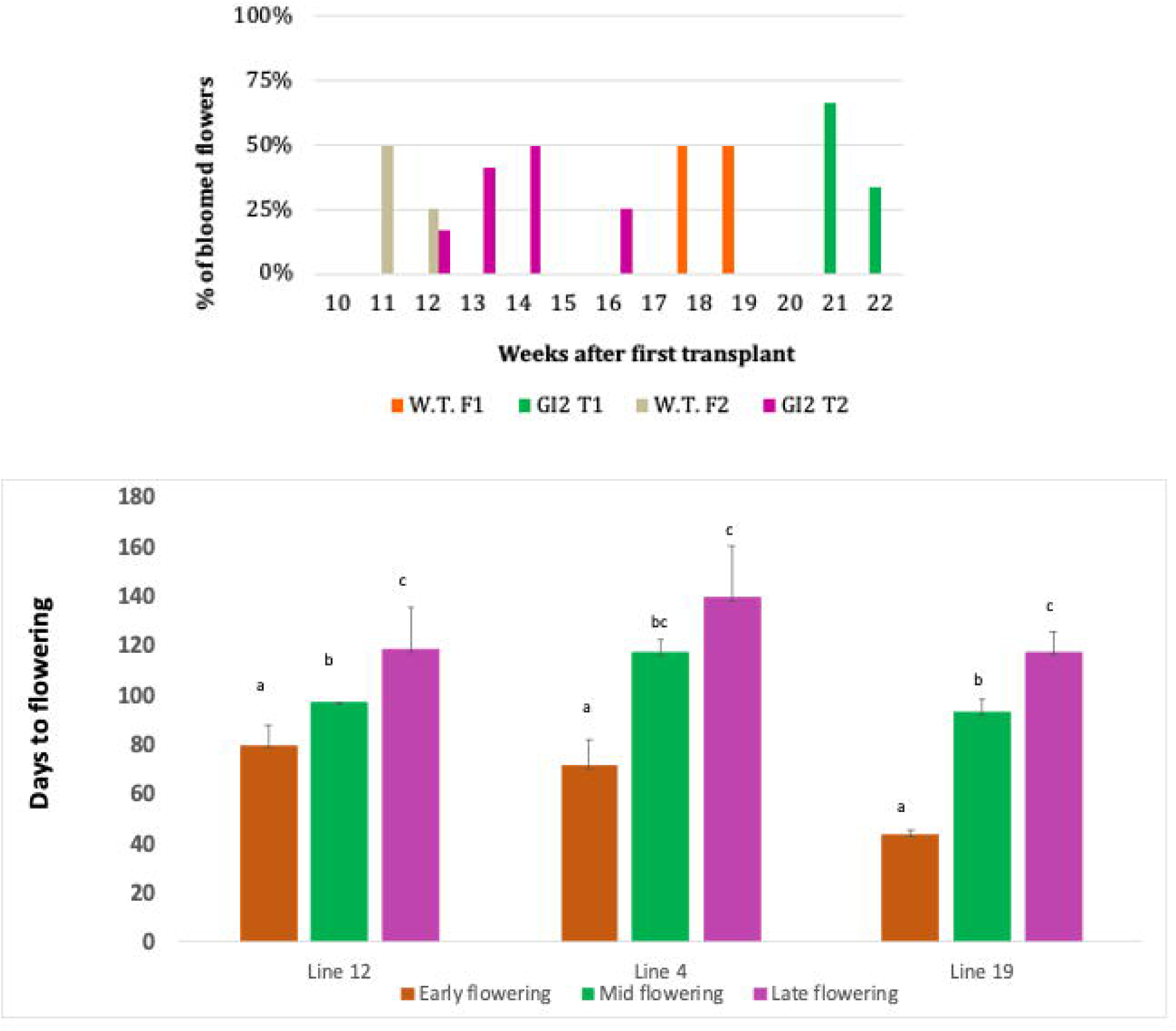
Effect of *PhGI2* downregulation and knockout on flowering time. (A) Percentage of fully open flowers in weeks after transplanting from *in vitro* culture to substrate of T1 and T2 generation of iRNA::*PhGI2* lines compared to wild type plant. (B) Segregation of flowering time from CRISPR/Cas9-GI2 lines 4, 12,and 19. Values are the average and deviation of 6 early flowering, mid flowering and late flowering plants. Letters indicate significant differences for each line according to Students T-test.

In RNAi::*PhGI2* plants, delayed flowering was accompanied by a reduction in the total number of flower buds at the end of the flowering period ranging between 32 and 86% compared to WT (Supplementary Information Table S4). Additionally, 38.5% of these buds emerged ectopically as a third lateral organ positioned between the terminal flower and the inflorescence shoot (Fig. 4A-D). Most ectopic flowers aborted prematurely, ultimately resulting in a reduction of fully developed flowers by 35% compared to wild-type controls. The corolla diameter was reduced on average by 31% and floral tube length by 13%. This suggested that *PhGI2* could have a similar role to *PhGI1* preventing floral abortion. Alternatively, the observed phenotypes were the result of cosuppression of *PhGI1*.

**Fig. 4.**
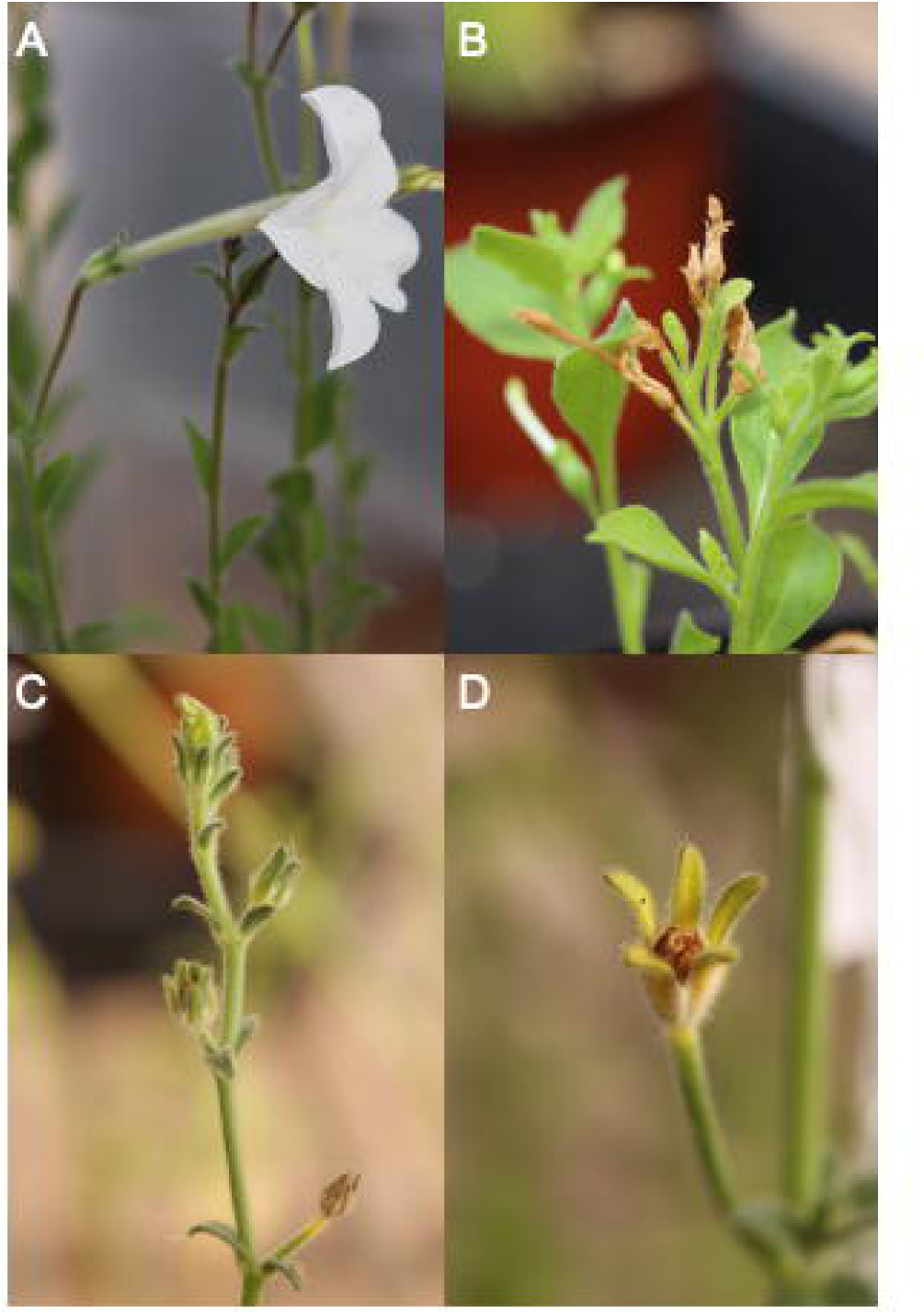
Effect of downregulation of *PhGI2* on flower development. (A) Wild type Petunia flower. (B) RNAi::*PhGI2* inflorescence showing ectopic flowers and floral abortion. (c) Inflorescence showing aborted flowers with distinct pedicel death (d) Close up of early aborted flower.

The previous assumption that *PhGI2* may be involved in suppressing early floral abortion was not confirmed. Indeed, no aborted or ectopic flowers were observed in any of the CRISPR/Cas9-derived alleles or their segregating populations (data not shown). Furthermore, corolla width and tube length were not affected (Supplementary Table S5). These findings suggest that the floral abnormalities are specifically associated with reduced *PhGI1* transcript levels due to RNA interference. These data effectively rule out a function of *PhGI2* on floral development. Furthermore, as three independent *PhGI2* alleles exhibited a clear dosage-dependent effect on flowering time, we conclude that *PhGI2* plays a central role in regulating flowering time in Petunia. This dosage sensitivity likely reflects a broader biological function of GI across plant species (see Discussion).

### *PhGI2* is not involved in vegetative growth

To determine whether *PhGI2* influences vegetative development, we analysed both RNAi-silenced lines and CRISPR/Cas9-induced mutants. As flowering time differed by months, we took all measures when the first flowers appeared. RNAi lines exhibited marked alterations in shoot growth as the main stem showed a significant reduction of 54.7% (p = 0.0059) (Fig. 5A, Supplementary Table S6). These architectural changes were associated with shortened basal, median and apical internodes. The number of leaves before the first flower was also significantly reduced by 46.7% (0=0.000).

**Fig. 5.**
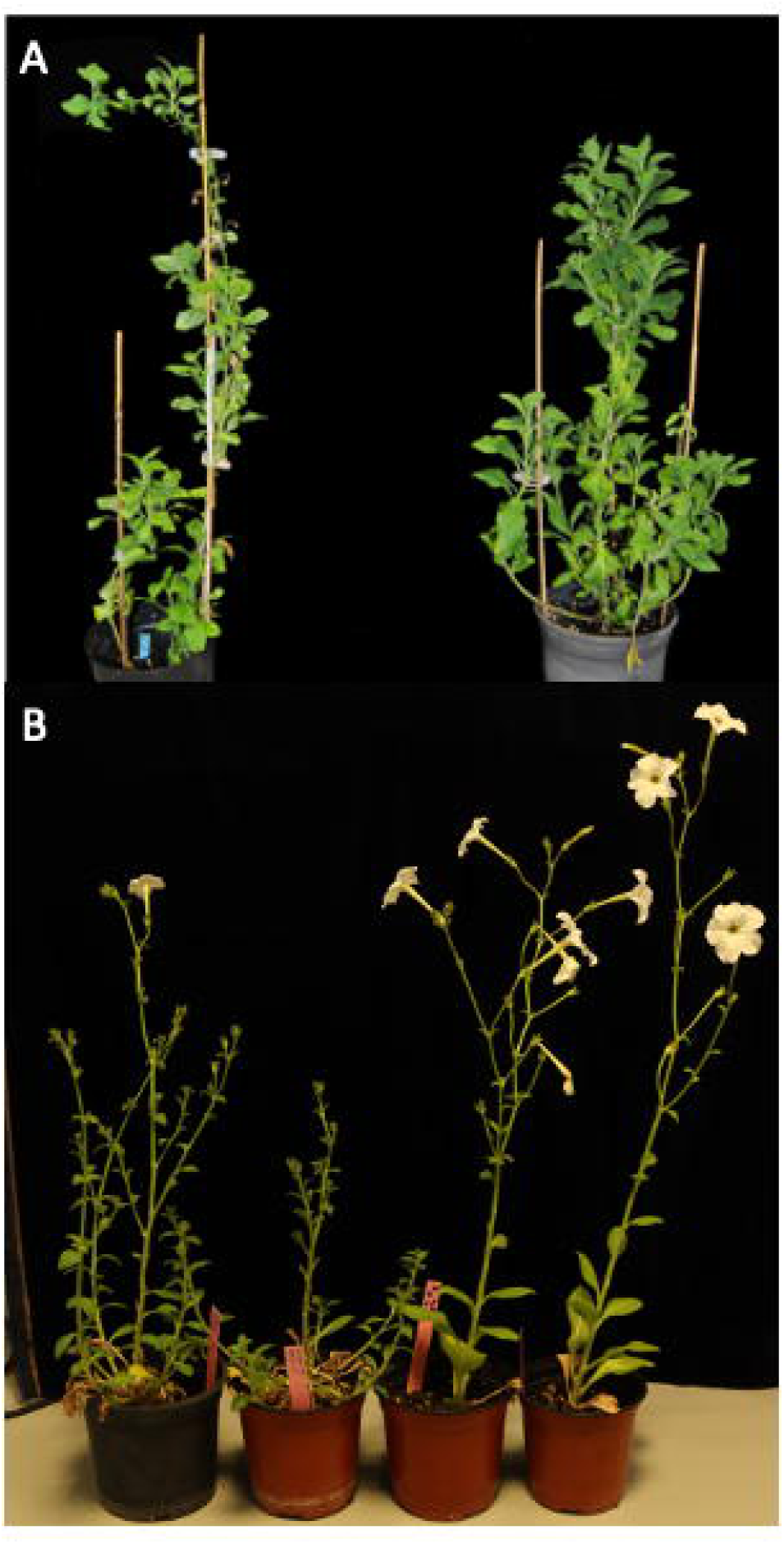
Effect of downregulation and knockout of *PhGI2* on vegetative development. (A) Vegetative growth characteristics in WT (left) and iRNA::*PhGI2* (right) plants under growth chamber conditions of 16 hours light/8 hours darkness. (B) From left to right, heterozygote, homozygous *PhGI2.12* and two WT plants grown in the greenhouse.

In contrast, *PhGI2* CRISPR mutants did not exhibit any significant changes in vegetative morphology. Despite the clear dosage-dependent effect on flowering time where WT plants flowered early, heterozygotes intermediate, and homozygous mutants late (Fig. 5 B), all late flowering genotypes were morphologically indistinguishable from each other. No differences were observed in plant height, branch number, internode length, or leaf size (Supplementary Information Table S7). These findings collectively indicate that *PhGI2* primarily regulates flowering time, and that vegetative development can be genetically uncoupled from floral induction.

### *PhGI2* is not involved in scent control

We measured the emission of VOCs from the flowers of four plants of the independent lines 4 and 6 of RNAi::*PhGI2*, and in wildtype. We distinguished main VOCs characterized by a threshold level of 2% of total VOCs (Table S8), and minor VOCs. *RNAi::PhG2* lines showed a reduction on average of 19.42% in total VOC emission (Fig. 6A). We analysed rhythmic VOC emission during 24 hours in 3-hour intervals (Fig. 6B). In silenced lines, lowest emission was recorded at ZT9, three hours later than in wildtype flowers. Both wildtype and silenced lines showed peak emission during the dark phase at ZT18.

**Fig. 6.**
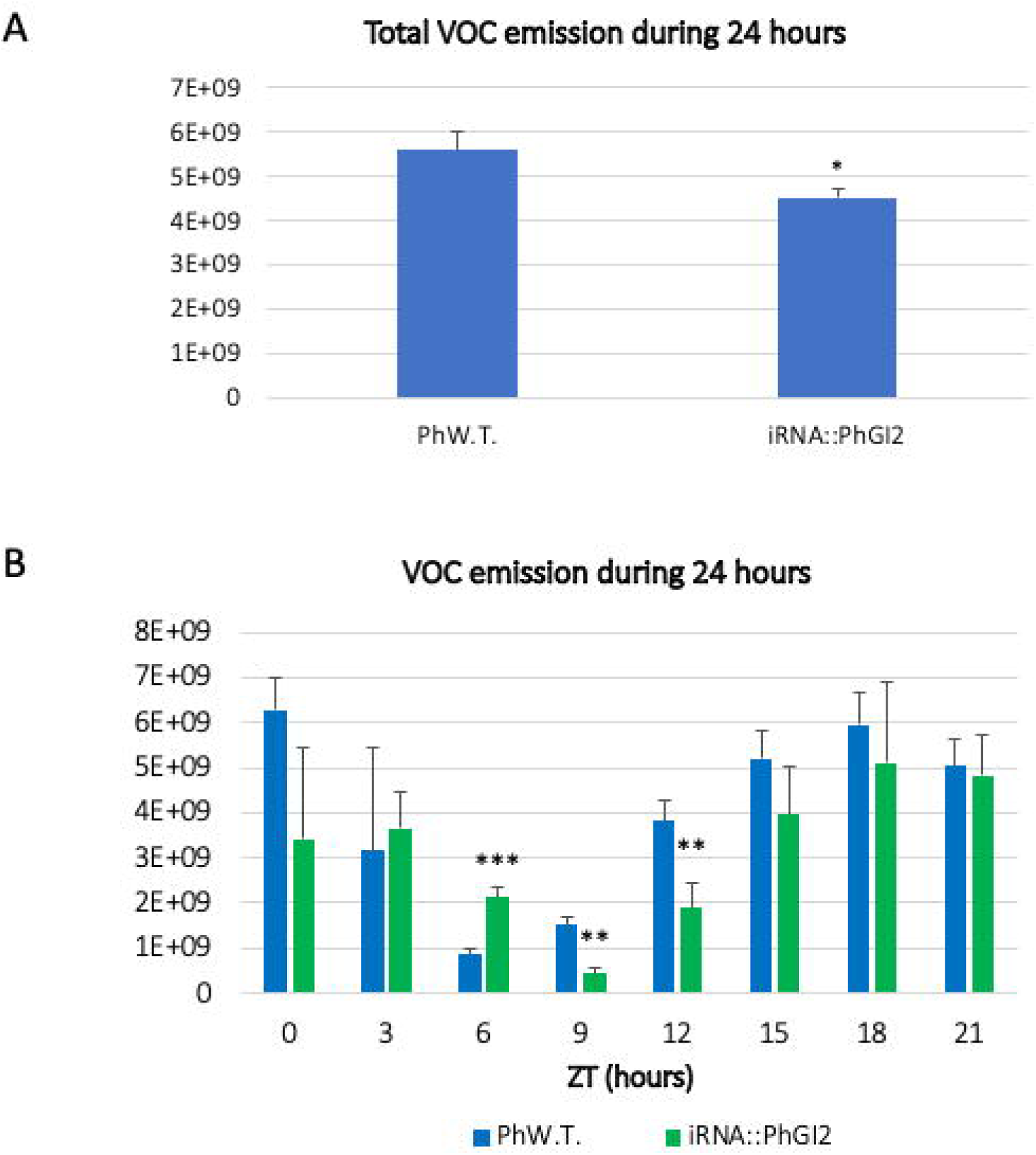
Volatile emission by flowers from wild type and iRNA::P*hGI2* T1 lines. Flowers were excised at ZT0. (A) Total VOC emission in wild type flowers compared to iRNA::*PhGI2* lines in 24 hours and (B) VOCs emission In three hour intervals during 24 hours. Absolute total emission of VOCs per grams of fresh weight is given as sum of integrated peak area. Asterisks indicate statistical significance between wild type and iRNA lines with *P < 0.05; **P < 0.01; ***P < 0.001 according to Student’s T-test.

We compared VOC emission between early flowering WT siblings to late flowering homozygous mutants of *PhGI2.4* and *PhGI2.12* in a 4-hour sampling period between ZT0-4. There were no significant differences between the samples in the amount of methyl benzoate, main and total VOCs (Fig. 7A). The VOC pattern of the main volatiles excluding methyl benzoate did not change consistently between late and early siblings or the WT (Fig. 7B). Thus, we conclude that *PhGI2* is not involved in coordinating quality or quantitative scent emission. The observed decrease in total VOC production and changes in rhythmic emission in RNAi lines are probably an effect of combined reduction of *PhGI1* and *PhGI2*.

**Fig. 7.**
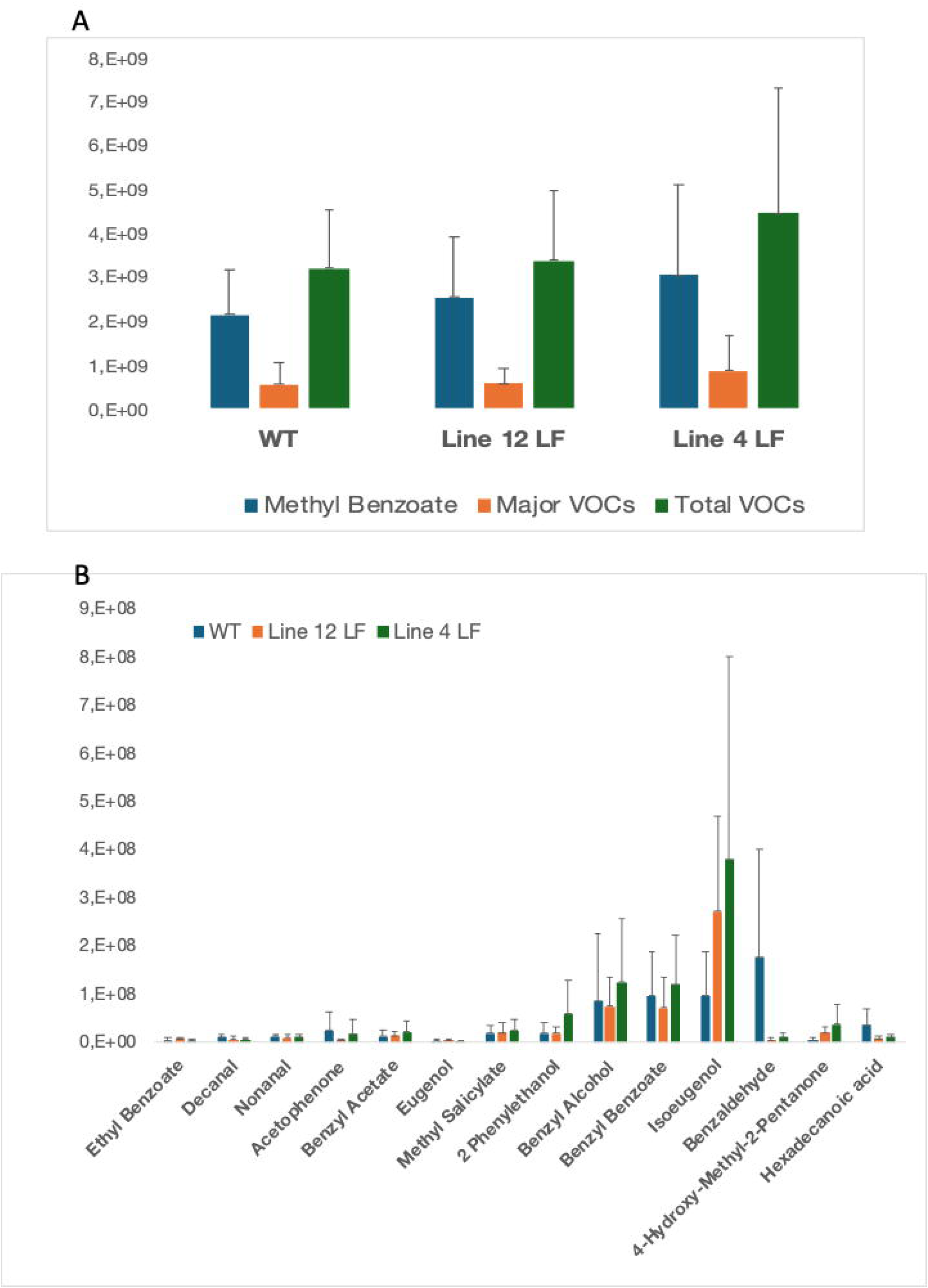
Effect of *PhGI2* knockout on VOC emission. (A) Amount of methyl benzoate, main VOCs and total VOCs and (B) amount of main VOCs of wild type flowers and homozygote *PhGI2.12* and *PhGI2.4* flowers. Emission was recorded during 4 hours from ZT 0 to ZT4. and calculated based on the integrated peak area divided by flower fresh weight.

## Discussion

In this work we have performed a comprehensive analysis of *PhGI2* and its biological functions. *GI* expression was originally elucidated in Arabidopsis where it shows a strong circadian oscillation. Most of the transcriptomic studies of circadian clock have been performed using leaves. While the exact peak of expression may not be conserved among plant species, *GI* is one of the clock genes with a cycling expression pattern (Izawa *et al*., 2011; Berns *et al*., 2014; Terry *et al*., 2019a). In wildtype plants, the expression pattern of *PhGI2* was characterized by an evening phased peak expression, coinciding with *PhGI1* expression and similar to the expression pattern observed for *GI* orthologs in other species, including Arabidopsis (Fowler *et al*., 1999), or soybean (Marcolino-Gomes *et al*., 2014). In contrast to Arabidopsis, where *AtGI* shows strong oscillation under free running conditions (Park *et al*., 1999), *PhGI2* cyclic expression in leaves depends on photoperiod and is identical to *PhGI1* (Fenske *et al*., 2015; Brandoli *et al*., 2020). Classic work showed that Arabidopsis hypocotyl elongation, leaf growth and movement are circadian regulated i.e. they are somehow resilient to free running conditions of continuous light or dark (Millar *et al*., 1995; Apelt *et al*., 2017). We had previously shown that petunia leaf movement depends on light inputs (Díaz-Galián *et al*., 2019). As *PhGI1* and *PhGI2* expression is also light dependent, we conclude that the Petunia circadian clock has a different configuration, both in terms of external cues and internal coordination.

Although we attempted to down regulate *PhGI2* in a specific manner, designing *RNAi* constructs targeting the 3’UTR, this was not achieved and *PhGI1* was readily down regulated to over 50%. Classic work in Petunia has shown that RNAi based downregulation may cause a strong cosuppression of genes with high DNA homology (Angenent *et al*., 1994; Jorgensen, 1995; Paoli *et al*., 2009). Our data indicates that the common phenotypes found in downregulated plants of *PhGI1* or *PhGI2* are in fact an effect of *PhGI1* silencing and not *PhGI2* as most of them are not found in the *PhGI2* CRISPR alleles identified.

We had previously hypothesized that *PhGI1* and *PhGI2* expression maybe mutually regulated. Indeed, the *PhGI2.12* mutant has a drastic effect on the amplitude and rhythmicity of *PhGI1*, *PhTOC, PhLHY* and its own expression. Considering the lack of data concerning *GI* as a transcriptional regulator, the molecular mechanism of this uncovered autoregulation needs further molecular analysis. Recent work suggests that *GI* may cooperate with transcription factors in controlling target genes, but the data available is correlational (Siemiatkowska *et al*., 2022).

Despite the dramatic shift in flowering time, vegetative growth remained unaltered in *PhGI2* mutants. This is different from *Arabidopsis*, where late-flowering genotypes, including *AtGI*, named for its vegetative phenotype, typically display increased biomass. The functions of *GI* in flowering occur via the *FT*-*TFL1* loci. Extensive research in day-neutral tomato and other species has shown that the fine-tuning of *FT* (downstream of *GI*) and *SP/TFL1/FD* regulates determinate growth (Lifschitz *et al*., 2006; Shalit *et al*., 2009). Supporting this model, the loss of function of *FT* and *TFL1* paralogs in *Petunia*, as well as a QTL in common bean (*PvTFL1*), has been shown to coordinate flowering and vegetative development (González *et al*., 2016; Abdulla *et al*., 2024). Importantly, vegetative growth has two layers. One is determinate versus indeterminate growth and the second, lateral organ size. CRISPR-generated lines *PhGI2.12*, *PhGI2.4*, and *PhGI2.19* reveal that in *Petunia × hybrida*, flowering time and lateral organ size can be genetically uncoupled. Although increased biomass may correlate with delayed flowering, it is not directly caused by it. Moreover, *PhGI1* appears to have undergone subfunctionalization to repress vegetative growth, while *PhGI2* retains a role in promoting floral transition. These findings support the idea that floral induction and reduced vegetative growth are parallel, rather than causally linked, developmental processes. They also highlight the two distinct aspects of vegetative growth, lateral organ size and growth termination, as genetically and functionally separate components.

It is worth noting that homozygous *PhGI2* CRISPR alleles may take up to 140 days vs 70 to flower, compared to segregating WT siblings. The *GI* locus is involved in control of floral transition in many other species. The quantitative function of *GI* appears to be conserved in a variety of plants as *OsGI* has a dosage effect in rice under field conditions (Izawa *et al*., 2011). Furthermore, in soybean *GmGI* has three paralogs and they are involved in domestication and adaptation to different environmental conditions (Wang *et al*., 2023). Our finding of a dosage effect of *PhGI2* on flowering time indicates that it is a major regulator of this trait across land plants.

In *Petunia x hybrida*, the emission of floral fragrance is dominated by benzenoids. The lack of consistent changes in VOC profiles in *PhGI2.4* and *PhGI2.12* suggest that other clock components may be involved in coordinating scent emission. Indeed work performed in *N.attenuata*, snapdragon and Petunia shows that *LHY* and *ZTL/CHL* are responsible for the circadian regulation of scent (Fenske *et al*., 2015; Yon *et al*., 2016; Terry *et al*., 2019b).

Taking all our data together we propose a model where the ancestral *GI* biological functions include repression of biomass, and induction of floral transition (Fig. 8A). The triple duplication of Solanaceae genomes gave rise to two paralogs in *Petunia x hybrida*, where *PhGI2* maintained the functions controlling floral transition and shared with *PhGI1* the circadian clock coordination. In contrast the repression of vegetative growth and all the newly uncovered functions in flower development resulted from a subfunctionalization and neofunctionalization of *PhGI1* (Fig. 8B). Coupling of floral transition and vegetative development in Arabidopsis may differ from Petunia in its regulatory coordination as Arabidopsis inflorescences are racemes while Petunia and other Solanaceae have cymose inflorescences (Prusinkiewicz *et al*., 2007; Rebocho *et al*., 2008). Our work describes for the first time the functional evolution of circadian clock genes in Solanaceae and shows a genetic separation of vegetative growth from flower transition.

**Fig. 8.**
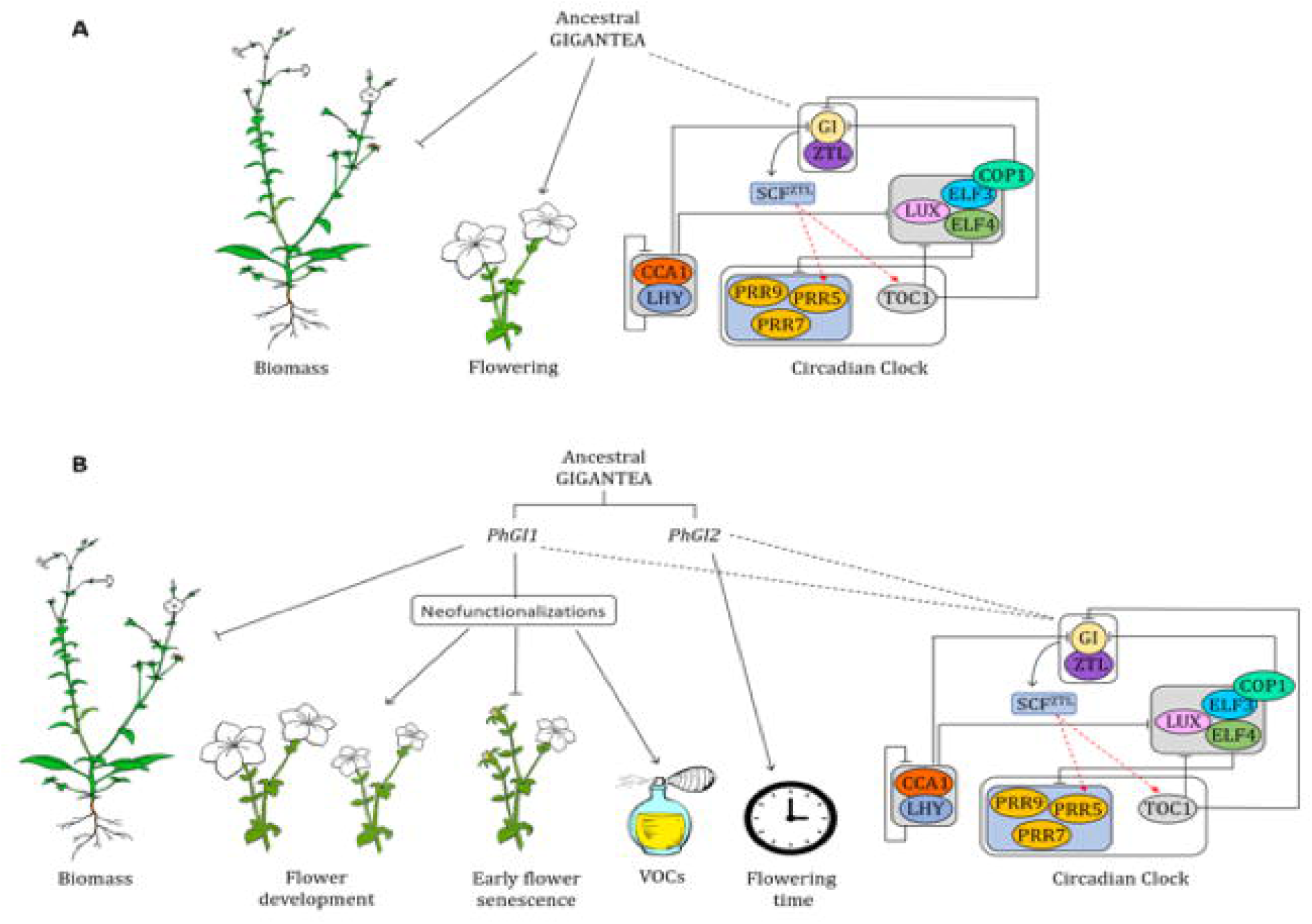
(A) Model for the ancestral *GI* gene function and (B) model for function of *GI* paralogs in Petunia after a genome triplication. *PhGI2* retained the floral induction and circadian functions while *PhGI1* retained growth repression and evolved new roles in flower development.

## Acknowledgments

The current research was funded by project PID2021-127933OB-C21 funded by Ministerio de Ciencia, Innovación y Universidades from Spain.

The authors declare that no generative AI tools were used to create scientific content, data, or interpretations in this manuscript. Generative AI (ChatGPT) was used exclusively for language editing to improve clarity and grammar. All authors reviewed and approved the final text.

## Competing Interests

None declared

## Author contributions

C.B, design of the research; performance of the research; data analysis, collection, interpretation, figure development, writing the manuscript; MEC design of the research; performance of the research; data analysis, interpretation, writing the manuscript, funding acquisition, project management; JW design of the research; performance of the research; data analysis, collection, interpretation, figure development, writing the manuscript, funding acquisition, project management.

## Data Availability

## Supplementary information

**Fig. S1** Multiple sequence alignment of GI proteins from land plants.

**Fig. S2** Expression of (a) PhGI2, (b) PhGI1, (c) PhZTL, (d) PhLHY, (e) PhTOC and (f) PhELF4 in iRNA::PhGI2 lines 4.4, 6.1 and 6.3 compared to expression in the wild-type, during a 24 hour period.

**Table S1** PCR primers.

**Table S2** Statistical analysis of rhythmicity of *PhGI1* and *PhGI2* gene expression data under normal (LD) and free running conditions (DD).

**Table S3.** Statistical analysis of rhythmicity of gene expression data in RNAi lines

**Table S4.** Statistical analysis of rhythmicity of gene expression data in *PhGI2*.12 mutant

**Table S5** Segregation of three CRISPR/Cas9 alleles of PhGI2 for flowering time.

**Table S6** Comparison of floral parameters between Wild-Type and silenced *PhGI2*.

**Table S7** Comparison of floral parameters between WT (early flowering). mid (heterozygous) and late (homozygous) flowering plants of Crispr/Cas9 *PhGI2* lines 12. 4 and 19.

**Table S8** Vegetative parameters of iRNA::*PhGI2* in T2 generation.

**Table S9** Comparison of vegetative parameters between late flowering Crispr/Cas9 *Gi2* mutants (late flowering) and early and medium flowering siblings.

**Table S10** Main Volatile Organic Compounds analyzed.

## Notes

### Competing Interest Statement

The authors have declared no competing interest.

